# Complete genome sequence of *Xylella taiwanensis* and comparative analysis of virulence gene content with *Xylella fastidiosa*

**DOI:** 10.1101/2021.03.08.434500

**Authors:** Ling-Wei Weng, Yu-Chen Lin, Chiou-Chu Su, Ching-Ting Huang, Shu-Ting Cho, Ai-Ping Chen, Shu-Jen Chou, Chi-Wei Tsai, Chih-Horng Kuo

## Abstract

The bacterial genus *Xylella* contains plant pathogens that are major threats to agriculture in America and Europe. Although extensive research was conducted to characterize different subspecies of *Xylella fastidiosa* (*Xf*), comparative analysis at above-species levels were lacking due to the unavailability of appropriate data sets. Recently, a bacterium that causes pear leaf scorch (PLS) in Taiwan was described as the second *Xylella* species (i.e., *Xylella taiwanensis*; *Xt*). In this work, we report the complete genome sequence of *Xt* type strain PLS229^T^. The genome-scale phylogeny provided strong support that *Xf* subspecies *pauca* (*Xfp*) is the basal lineage of this species and *Xylella* was derived from the paraphyletic genus *Xanthomonas*. Quantification of genomic divergence indicated that different *Xf* subspecies share ∼87-95% of their chromosomal segments, while the two *Xylella* species share only ∼66-70%. Analysis of overall gene content suggested that *Xt* is most similar to *Xf* subspecies *sandyi* (*Xfs*). Based on the existing knowledge of *Xf* virulence genes, the homolog distribution among 28 *Xylella* representatives was examined. Among the 11 functional categories, those involved in secretion and metabolism are the most conserved ones with no copy number variation. In contrast, several genes related to adhesins, hydrolytic enzymes, and toxin-antitoxin systems are highly variable in their copy numbers. Those virulence genes with high levels of conservation or variation may be promising candidates for future studies. In summary, the new genome sequence and analysis reported in this work contributed to the study of several important pathogens in the family Xanthomonadaceae.

**Contribution to the Field:** *Xylella fastidiosa* is a plant-pathogenic bacterium with multiple subspecies that are major threats to agriculture in America and Europe. Although extensive research has been conducted, comparative analysis of this species with other bacteria is lacking due to the unavailability of known close relatives. In this work, we report the complete genome sequence of *Xylella taiwanensis*, a newly described species within the same genus. This new data set and our focused analysis helped to better understand the evolutionary relationships among different *Xylella* lineages and their genomic diversity. Moreover, detailed examination of their virulence genes identified those that are either highly conserved or variable, providing promising candidates for future studies to further investigate the molecular mechanisms of *Xylella* virulence.

## Introduction

The gammaproteobacterium *Xylella fastidiosa* (*Xf*) is an insect-vectored plant pathogen that resides in plant xylem and is fastidious (Wells et al., 1987). To date, more than 563 plant species in 82 families have been reported as hosts for *Xf* (European Food Safety Authority, 2018). *Xf* could be classified into at least five subspecies; some of the notable examples include *Xf* subspecies *fastidiosa* (*Xff*) that causes Pierce’s disease (PD) of grapevine, *Xf* subspecies *pauca* (*Xfp*) that causes citrus variegated chlorosis (CVC) and olive quick decline syndrome (OQDS), and *Xf* subspecies *sandyi* (*Xfs*) that causes oleander leaf scorch (OLS). Because of their economic and ecological impacts, substantial resources have been devoted to related research. Notably, several large-scale studies were conducted to investigate the genomic diversity and evolution of *Xf* (Denancé et al., 2019; Potnis et al., 2019; Vanhove et al., 2019). Based on a comparison of 72 strains, the five *Xf* subspecies harbor high levels of genetic diversity (Vanhove et al., 2019). With an average gene content of ∼2,150 per strain, the core genome (i.e., genes shared by >95% of the strains) contains only ∼900 genes, while the pangenome contains ∼10,000 genes. Moreover, although certain patterns of sequence divergence were found among those subspecies (Denancé et al., 2019), extensive recombination occurred at the levels of within- and between-subspecies (Potnis et al., 2019).

In contrast to the extensive genomic research at within-species level, comparative studies of *Xf* at higher taxonomic levels are lacking. Under the current taxonomy, *Xylella* belongs to the family Xanthomonadaceae and is most closely related to *Xanthomonas* (Rodriguez-R et al., 2012; Anderson et al., 2013). However, the genomic divergence between *Xylella* and *Xanthomonas* is very high in terms of chromosomal organization, gene content, and sequence variation. Thus, extracting biological insights from such comparisons is difficult. At within-genus level, *Xf* was largely considered as the only species within this genus since it was formally described in 1987 (Wells et al., 1987), which made between-species comparison infeasible. Intriguingly, a *Xylella* lineage that causes pear leaf scorch (PLS) in Taiwan was found to exhibit a slightly lower level of 16S rRNA gene sequence identity at 97.8-98.6% when compared to different subspecies of *Xf* (Su et al., 2012). In 2016, this PLS *Xylella* was formally reclassified as a novel species *Xylella taiwanensis* (*Xt*) based on a polyphasic approach (Su et al., 2016). Although a draft genome sequence of *Xt* was published earlier (Su et al., 2014), that draft assembly was produced with only ∼20-fold coverage of Roche/454 GS-FLX reads and is highly fragmented (i.e., 85 contigs; N50 = 121 kb). Moreover, no comparative analysis of gene content between *Xt* and *Xf* has been conducted.

To fill this gap, we determined the complete genome sequence of the type strain of *Xt* (i.e., PLS229^T^) for comparative analysis with its relatives. In addition to providing a genome-level overview of their diversity and evolution, we utilized the existing knowledge of *Xff* virulence genes and conducted detailed comparisons of virulence gene content among different *Xylella* lineages.

## Materials and Methods

The strain was acquired from the Bioresource Collection and Research Centre (BCRC) in Taiwan (accession 80915). The procedures for genome sequencing and comparative analysis were based on those described in our previous studies (Lo et al., 2013; Lo et al., 2018; Cho et al., 2020). All bioinformatics tools were used with the default settings unless stated otherwise.

Briefly, the strain was cultivated on PD2 medium as described (Su et al., 2016) for DNA extraction using Wizard Genomic DNA Purification Kit (A1120; Promega, USA). For Illumina sequencing, a paired-end library with a target insert size of 550-bp was prepared using KAPA LTP Library Preparation Kit (KK8232; Roche, Switzerland) without amplification, then sequenced using MiSeq Reagent Nano Kit v2 (MS-103-1003; Illumina, USA) to obtain ∼50X coverage. For Oxford Nanopore Technologies (ONT) sequencing, the library was prepared using ONT Ligation Kit (SQK-LSK109) and sequenced using MinION (FLO-MIN106; R9.4 chemistry and MinKNOW Core v3.6.0) to obtain ∼228X coverage; Guppy v3.4.5 was used for basecalling. The raw reads were combined for *de novo* assembly by using Unicycler v0.4.8-beta (Wick et al., 2017). For validation, the Illumina and ONT raw reads were mapped to the assembly using BWA v0.7.12 (Li and Durbin, 2009) and Minimap2 v2.15 (Li, 2018), respectively. The results were programmatically checked using SAMtools v1.2 (Li et al., 2009) and manually inspected using IGV v2.3.57 (Robinson et al., 2011). The finalized assembly was submitted to the National Center for Biotechnology Information (NCBI) and annotated using their Prokaryotic Genome Annotation Pipeline (PGAP) (Tatusova et al., 2016).

A total of 40 genomes, including 27 from *Xf*, were used for comparative analysis (**Table 1**). Our taxon sampling mainly focused on the strains that could represent the known *Xylella* diversity (Vanhove et al., 2019). Two other Xanthomonadaceae genera were also included. For the closely-related *Xanthomonas*, 10 species were selected to represent the key lineages (Parkinson et al., 2009; Rodriguez-R et al., 2012). For the distantly-related *Pseudoxanthomonas*, only two species were sampled.

**Table 1.**
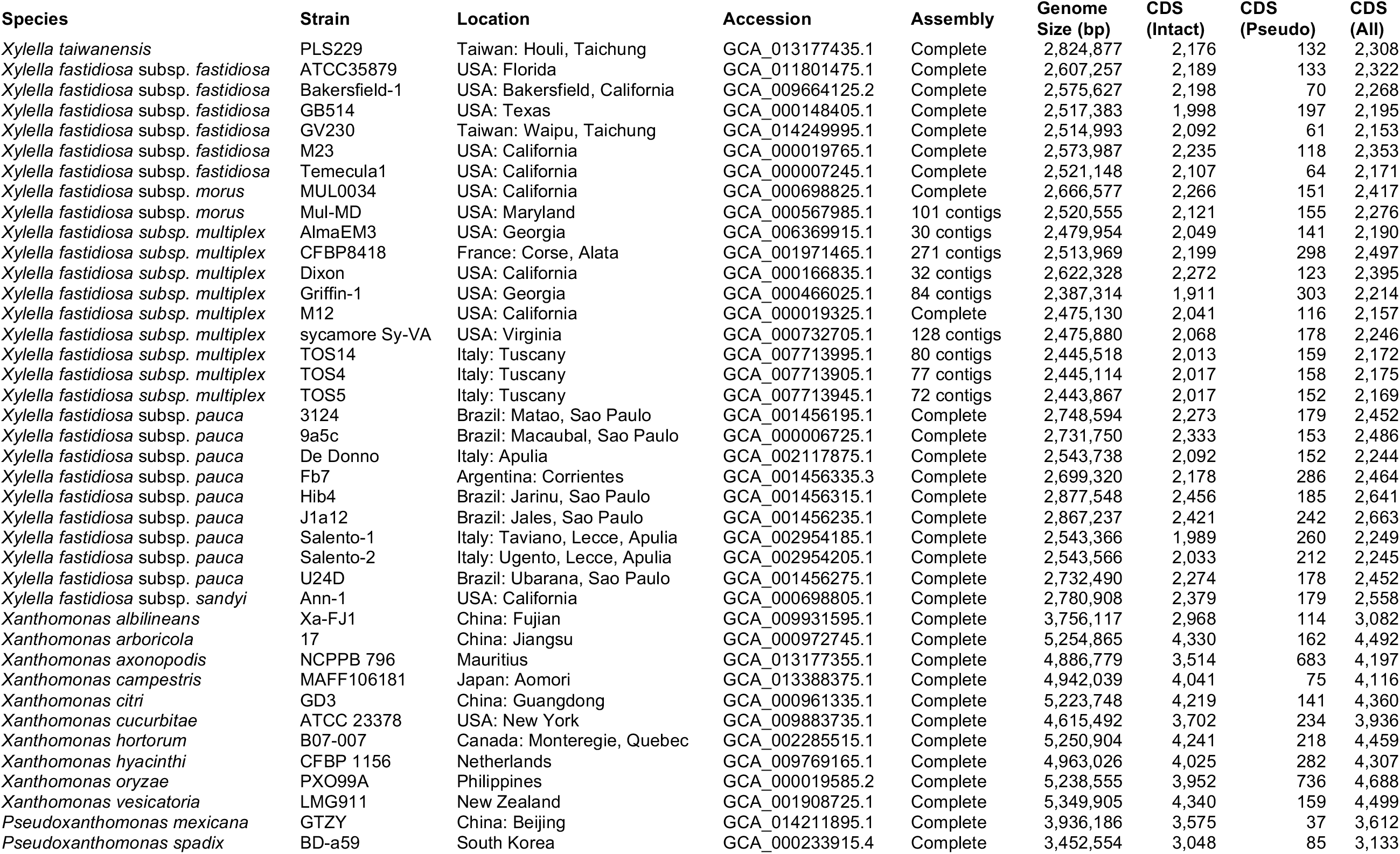
List of the genome sequences analyzed.

Chromosomal level comparisons of nucleotide sequences were conducted using fastANI v1.1 (Jain et al., 2018). Homologous gene clusters were identified based on protein sequences using BLASTP v2.10.0+ (Camacho et al., 2009) and OrthoMCL v1.3 (Li et al., 2003). For gene content comparisons, the homolog clustering result was converted into a matrix of genomes by homolog clusters with the value in each cell corresponding to the copy number. This matrix was converted into a Jaccard distance matrix among genomes using VEGAN package v2.5-6 in R, then processed using the principal coordinates analysis function in APE v5.4 (Paradis and Schliep, 2019) and visualized using ggplot2 v3.3.2 (Wickham, 2016). For phylogenetic analysis, homologous sequences were aligned using MUSCLE v3.8.31 (Edgar, 2004). The maximum likelihood inference was performed using PhyML v.3.3.20180214 (Guindon and Gascuel, 2003); the proportion of invariable sites and the gamma distribution parameter were estimated from the data set and the number of substitute rate categories was set to four. The PROTDIST program of PHYLIP v3.697 (Felsenstein, 1989) was used to calculate sequence similarities.

## Results and Discussion

### Genome Characteristics

Strain *Xt* PLS229^T^ has one 2,824,877-bp circular chromosome with 53.3% G+C content; no plasmid was found. The annotation contains two complete sets of 16S-23S-5S rRNA genes, 49 tRNA genes, and 2,176 intact coding sequences (CDSs). This genome size is near the upper range of those *Xf* representatives (median: 2.54 Mb; range: 2.39-2.88 Mb) and much smaller compared to *Xanthomonas* spp. (median: 5.09 Mb; range: 3.76-5.35 Mb) (**Table 1**). Among all 40 representative Xanthomonadaceae genomes, the genome sizes and the numbers of intact CDSs have a correlation coefficient of 0.989 (*p* < 2.2e^-16^). Compared to those *Xf* representatives with similar genome sizes (i.e., ∼2.73-2.88 Mb), such as those five *Xfp* strains from Brazil or the *Xfs* strain from the USA, the *Xt* PLS229^T^ genome has fewer intact CDSs (i.e., 2,273-2,456 vs. 2,173) and fewer pseudogenes (i.e., 153-242 vs. 132). It is unclear if these observations were caused by annotation artifacts or have biological meaning.

### Molecular Phylogeny and Genome Divergence

A total of 779 single-copy protein-coding genes were found to be shared by the 40 Xanthomonadaceae genomes compared (**Table 1**). Based on the concatenated alignment of the protein sequences derived from these genes, a robust maximum likelihood phylogeny was inferred (**Figure 1**). The availability of this *Xt* genome sequence provided a more appropriate outgroup to root the *Xf* phylogeny and further supported that *Xfp* is the basal lineage (Denancé et al., 2019; Potnis et al., 2019; Vanhove et al., 2019).

**Figure 1.**
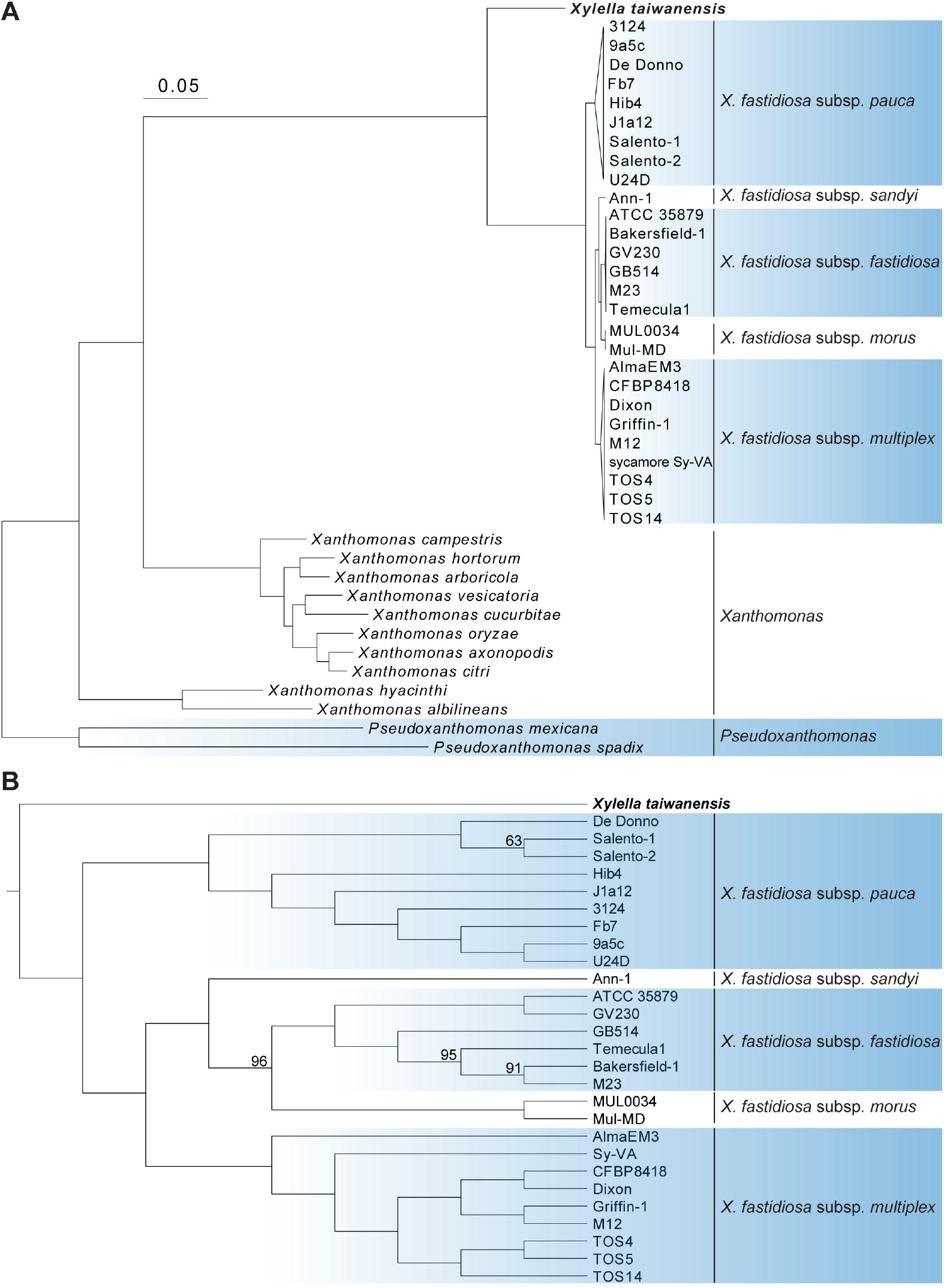
Molecular phylogeny of *Xylella* and related species in the family Xanthomonadaceae. The maximum likelihood phylogeny was based on 779 shared single-copy genes, the concatenated alignment contains 252,319 aligned amino acid sites. The genus *Pseudoxanthomonas* was included as the outgroup. (A) A phylogram for illustrating the relationships among all 40 genomes analyzed. Each of the *X. fastidiosa* subspecies was collapsed into a triangle for simplified visualization. All internal nodes illustrated in this phylogram received >95% bootstrap support based on 1,000 replicates. (B) A cladogram for illustrating the relationships among those 28 *Xylella* genomes analyzed. Internal nodes with bootstrap values lower than 100% were labeled.

The genus *Xanthomonas* was known to be paraphyletic but the relationships of its two major clades (i.e., represented by *Xanthomonas albilineans* and *Xanthomonas campestris*, respectively) with *Xylella* were controversial (Pieretti et al., 2009; Rodriguez-R et al., 2012). With our genome-scale phylogeny, it is clear that *Xylella* is more closely related to *X. campestris* (**Figure 1**) and has experienced genome reduction since their divergence (**Table 1**).

When the genetic divergence was measured by overall nucleotide sequence conservation using fastANI (Jain et al., 2018), comparisons within each of the five *Xf* subspecies found that 88.8-99.8% of the chromosomal segments are shared and those segments have 98.5-100% average nucleotide identity (ANI) (**Figure 2**). For between-subspecies comparisons, 86.6-94.8% of the chromosomal segments are shared and those segments have 96.3-98.8% ANI. When those *Xf* subspecies were compared to *Xt*, only 66.4-70.3% of the chromosomal segments are shared and those segments have 82.9-83.4% ANI. These results are consistent with previous findings (Su et al., 2016; Denancé et al., 2019) and provide further support to the current taxonomy based on the 95% ANI threshold recommended for delineating bacterial species (Jain et al., 2018).

**Figure 2.**
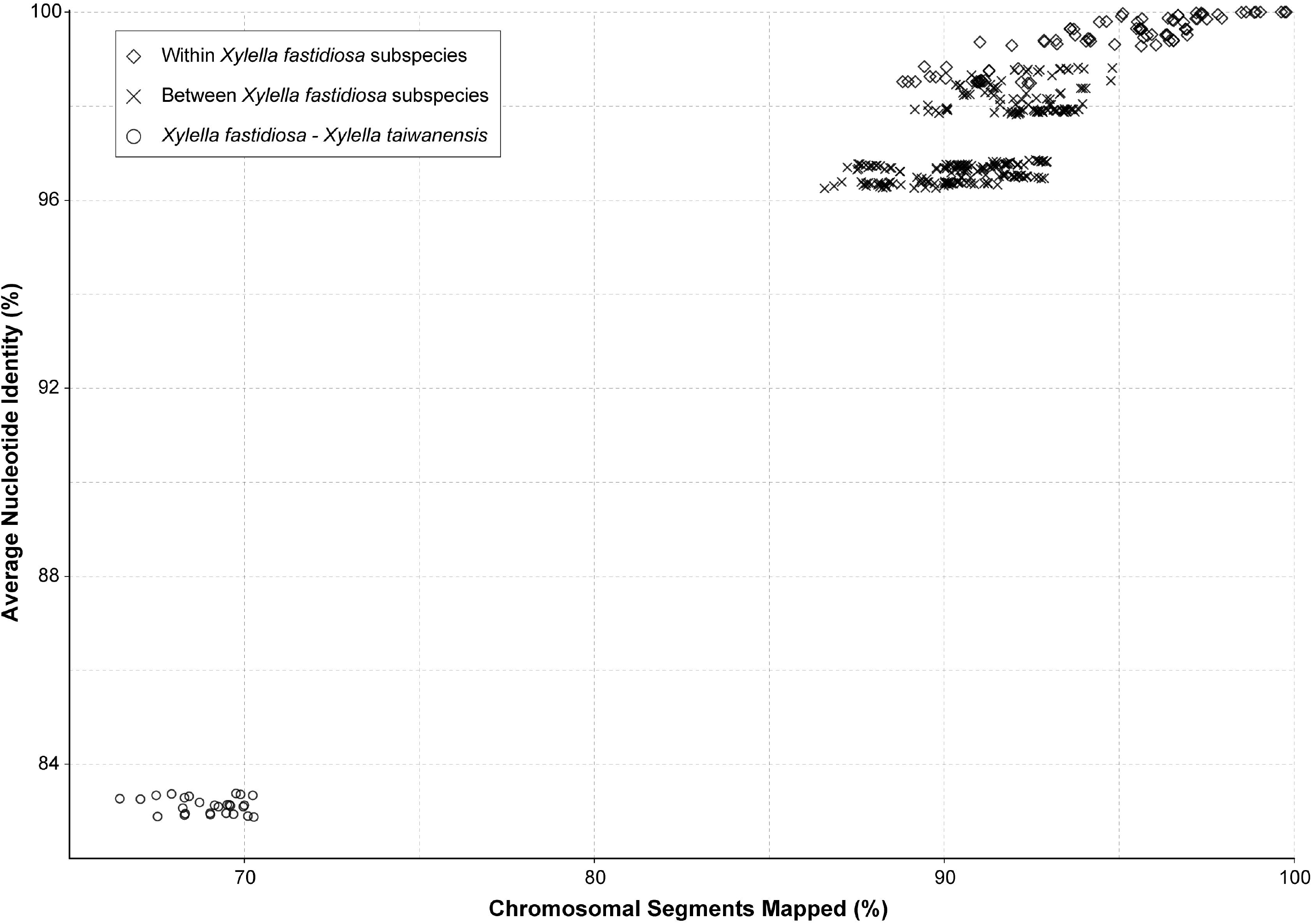
Genome similarity among the representative *Xylella* strains. The pairwise comparisons were classified into three categories: (1) within the same *X. fastidiosa* subspecies, (2) between different *X. fastidiosa* subspecies, and (3) between *X. fastidiosa* and *X. taiwanensis*.

Because the ANI approach provides low resolutions when the nucleotide sequence identity drops to ∼80% (Jain et al., 2018) and may not be appropriate for cross-genus comparisons, we also evaluated divergence based on the protein sequences of those 779 Xanthomonadaceae core genes. The two *Xylella* species have ∼88.8-89.1% protein sequence similarity, which is lower than the values observed in the comparisons among those eight *X. campestris* clade representatives (median: 93.8%; range: 92.6-97.2%), comparable to the *X. albilineans*-*X. hyacinthi* comparison (88.6%), and higher than the *P. mexicana*-*P. spadix* comparison (75.4%).

In addition to analysis of sequence divergence based on those 779 core genes (**Figure 1**) and the entire chromosomes (**Figure 2**), the divergence in gene content was also examined. The gene content comparisons were based on principal coordinates analysis that examines copy number variation among all homologous gene clusters in the entire pangenome and does not consider sequence divergence within each homologous gene cluster. When all 40 Xanthomonadaceae genomes were compared together based on their 11,455 homologous gene clusters, the grouping patterns (**Figure 3A**) are consistent with the phylogenetic clades inferred based on sequence divergence of the 779 core genes (**Figure 1A**). All 27 *Xf* genomes form a tight cluster (**Figure 3A**) despite their differences in the number of intact CDSs (range:1,911-2,456; av.± std. dev.: 2,156 ± 144) (**Table 1**). In contrast, although the *Xt* genome has 2,176 intact CDSs, which is close to the average observed among those 27 *Xf* representatives, it does not fall into the *Xf* cluster (**Figure 3A**). This result indicates that the gene content divergence between these two *Xylella* species is much higher than the divergence among *Xf* subspecies. For the within-*Xylella* comparison based on 5,395 homologous gene clusters, the grouping patterns are consistent with the taxonomic assignments and *Xt* is most similar to *Xfs* (**Figure 3B**). It is interesting that *Xt* and *Xfs* are similar in having a narrow host range (i.e., *Xt* is restricted to pear and *Xfs* is mostly known for oleander infections), while other *Xf* subspecies can infect a wide range of hosts (Baldi and Porta, 2017; European Food Safety Authority, 2018; Rapicavoli et al., 2018). However, it is also important to note that the host range information may be limited by sampling and experimental efforts. As more research results become available, this information may be updated. For example, *Xfs*-related strains have been reported to infect coffee (Jacques et al., 2016) and whether *Xt* can infect a wider range of plants remains to be investigated.

**Figure 3.**
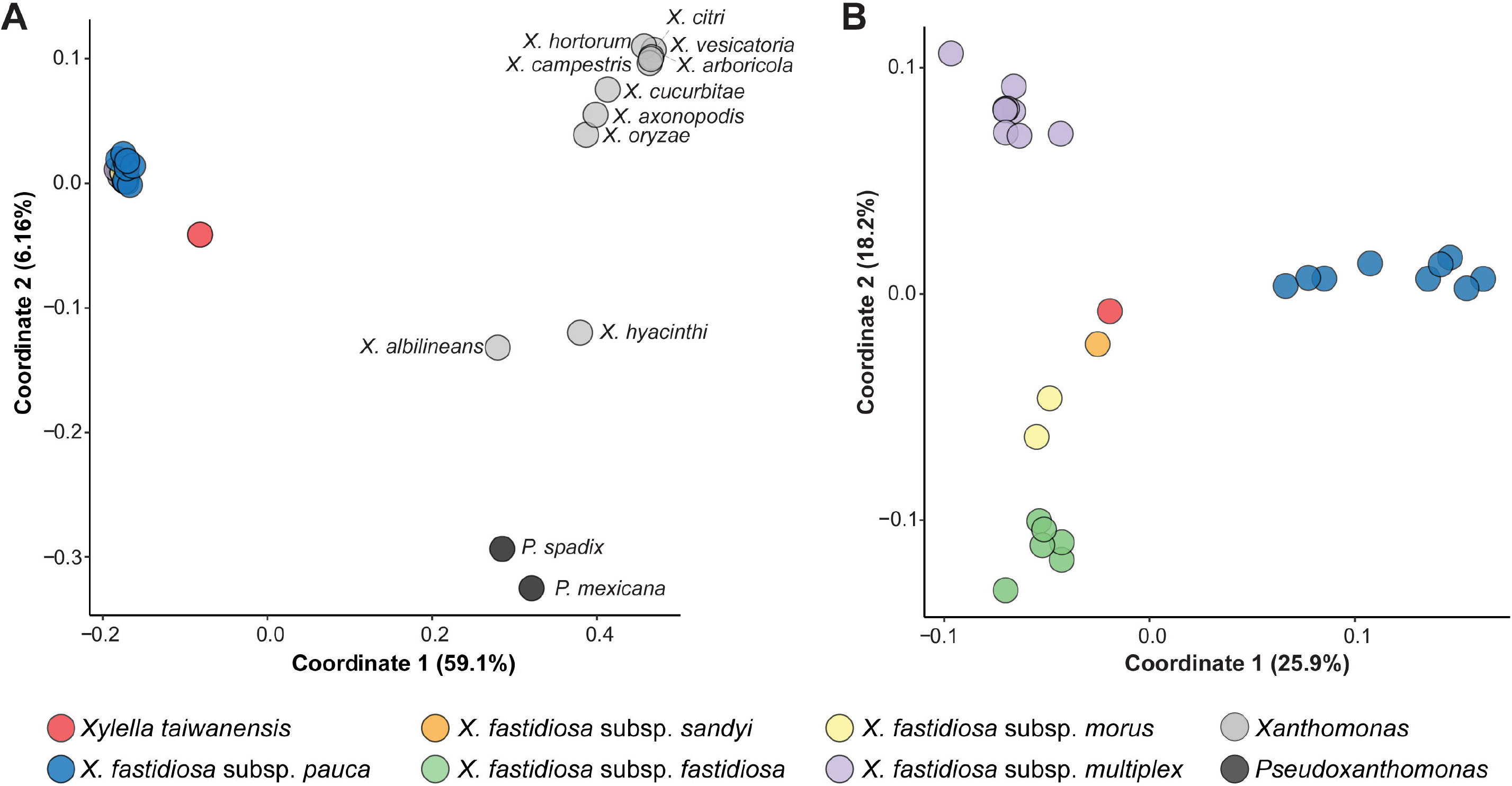
Principal coordinates analysis of gene content dissimilarity. The % variance explained by each coordinate was provided in parentheses. (A) Based on the 11,455 homologous gene clusters found among all 40 Xanthomonadaceae genomes analyzed in this work. (B) Based on the 5,395 homologous gene clusters found among the 28 *Xylella* genomes analyzed in this work.

### Virulence Genes and Pathogenicity Factors

Based on the current knowledge of putative virulence genes and pathogenicity factors identified in *Xf*, type IV pili (T4P) are important for twitching motility and movements within infected plants (Meng et al., 2005; Li et al., 2007; Burdman et al., 2011; Cursino et al., 2011; Hao et al., 2017). The *Xylella* T4P genes are organized into four major gene clusters with 25 homologs and highly conserved among the 28 representative genomes examined (**Figure 4**). Among these four T4P gene clusters, cluster II that corresponds to the *pil*-*chp* operon (Cursino et al., 2011; Hao et al., 2017) is the most conserved cluster with only two putative gene losses (i.e., *chpB* in *Xfp* Salento-1 and *chpC* in *Xt*). A notable gene absence is PD1925 in cluster IV, which encodes a hypothetical protein and is absent in all *Xfp* strains and *Xt*. The only gene duplication observed involves a tandem duplication of the cluster IV *pilA2* homolog in *Xt*. Intriguingly, *Xt* PLS229^T^ is the only strain that has two copies of *pilA2* and no *pilA1*. These two type IV pilin paralogs were shown to have different functions in *Xff*, with *pilA1* affecting pilus number and location while *pilA2* is required for twitching (Kandel et al., 2018). It is unclear if this *pilA2* duplication in *Xt* PLS229^T^ can complement its lack of *pilA1*.

**Figure 4.**
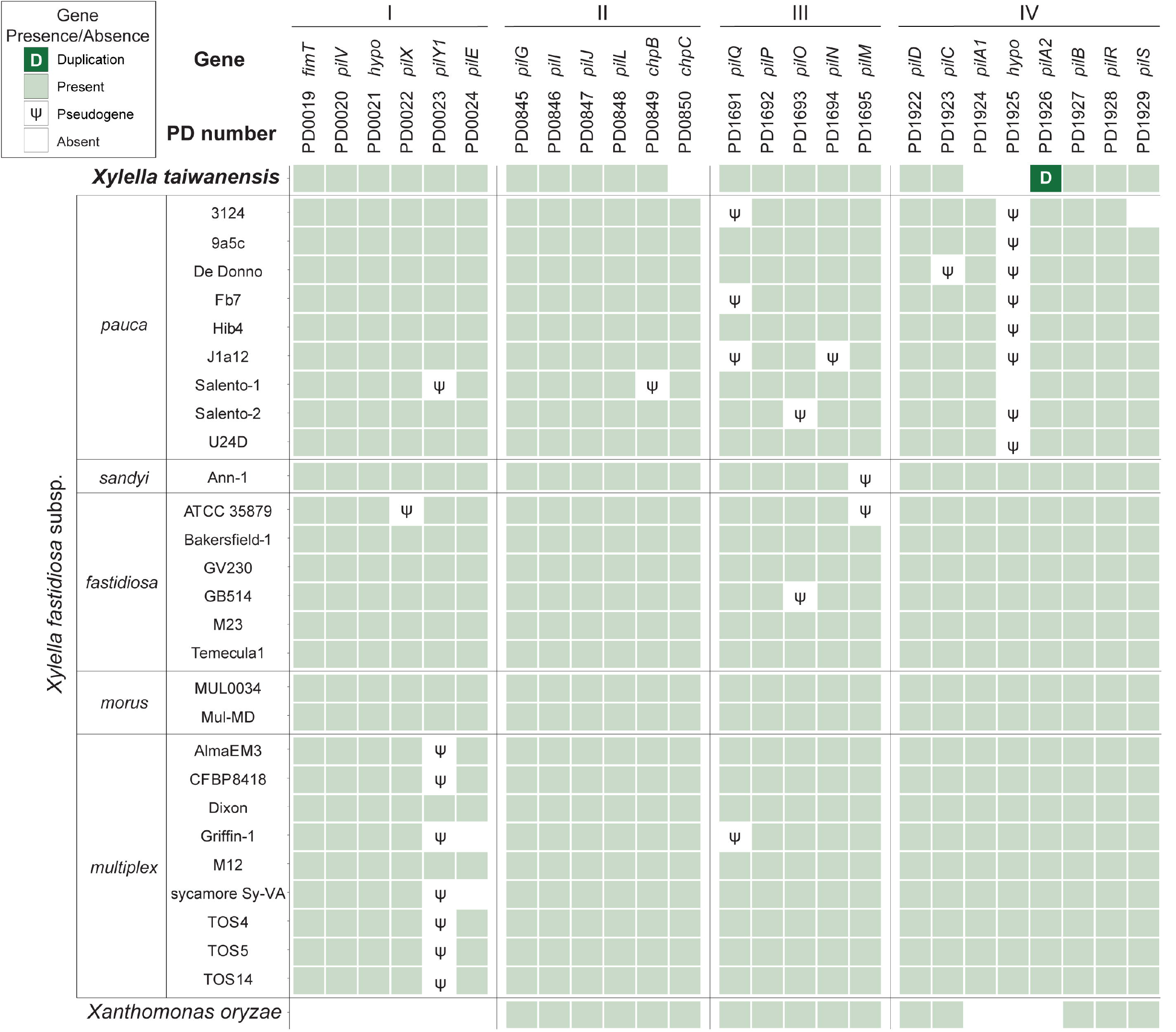
Distribution of the type IV pili genes found among representative *Xylella* genomes. *Xanthomonas oryzae* was included as the outgroup. The genes were identified by the PD numbers based on the annotation of Temecula1 genome; those with adjacent PD numbers are located in neighboring regions on the chromosome. A total of four gene clusters (labelled as I-IV) located in syntenic regions were found to be conserved. Gene names were provided when available; ‘*hypo*’ indicates those annotated as encoding hypothetical proteins. Patterns of gene presence and absence were illustrated in the format of a heatmap. For gene absence, those with identifiable pseudogenes were labelled accordingly. One case of tandem gene duplication was observed for the PD_1926 (*pilA2*) homolog in the *Xylella taiwanensis* genome.

In addition to the T4P genes, many other *Xylella* pathogenicity factors have been identified and these additional virulence genes may be classified into 10 major functional categories (Lee et al., 2014; Merfa et al., 2016; Chen et al., 2017; Rapicavoli et al., 2018; Ge et al., 2021). Among these categories, secretion systems and metabolism are the most conserved ones with no variation in gene copy number across all *Xylella* representatives (**Figure 5**). Additionally, those genes involved in regulatory systems are also highly conserved. In contrast, several genes related to adhesins, hydrolytic enzymes, and toxin-antitoxin systems are highly variable in copy numbers.

**Figure 5.**
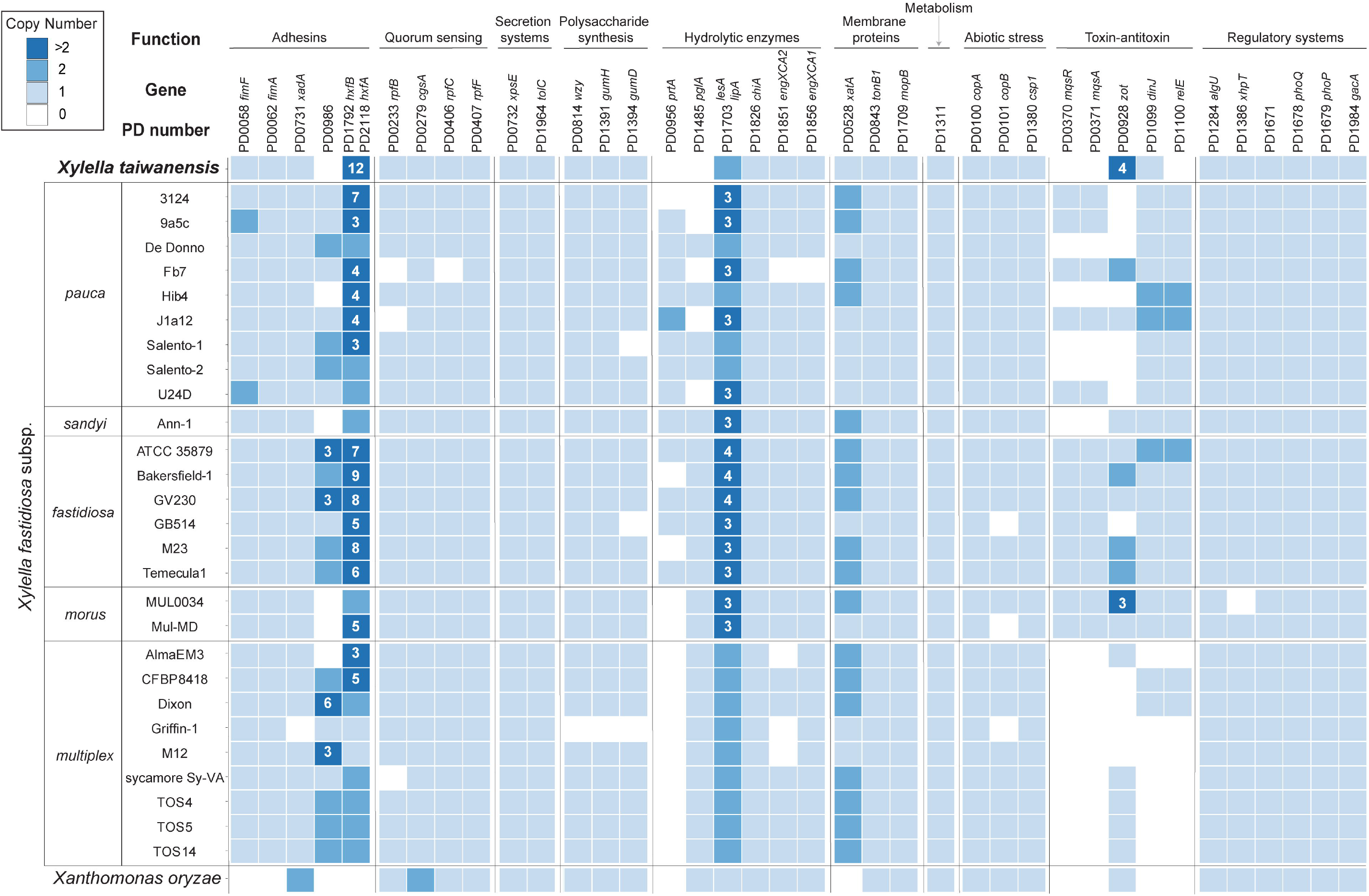
Distribution of putative virulence genes and pathogenicity factors among representative *Xylella* genomes. *Xanthomonas oryzae* was included as the outgroup. The homologous gene clusters were identified by the PD numbers based on the annotation of Temecula1 genome; gene names were provided when available. Gene copy numbers were illustrated in the format of a heatmap; values higher than two were labeled with the exact numbers. Two adhesin genes (i.e., PD1792 and PD2118) were assigned to the same homologous gene cluster and were combined for copy number calculation.

For more detailed examination, these putative virulence genes were classified into 38 homologous gene clusters and six are absent in the *Xt* genome (**Figure 5**). These include the genes that encode a putative adhesin (PD0986, hemagglutinin-like protein), two hydrolytic enzymes (PD0956, serine protease; PD1485, polygalacturonase), one pair of toxin-antitoxin (PD0370, motility quorum sensing regulator MqsR ribonuclease; PD0371, MqsA antitoxin), and another separate toxin (PD1100, endoribonuclease). Notably, the *mqsR*-*mqsA* toxin-antitoxin system genes (Lee et al., 2014; Merfa et al., 2016) are differentially distributed among those *Xf* subspecies. Homologs of these two genes are entirely conserved in all *Xff* and *Xf* subspecies *morus* strains, present in five out of the nine *Xfp* strains, and completely absent in *Xfs* and *Xf* subspecies *multiplex*.

Based on previous studies that characterized mutant phenotypes, PD0956 (Gouran et al., 2016) and PD1100 (Burbank and Stenger, 2017) are both antivirulence factors and the loss of either one resulted in hypervirulence of *Xff* in grapevines. Similarly, PD0370 is another antivirulence factor that reduces the virulence of *Xfp* against citrus when overexpressed (Merfa et al., 2016). In contrast, both PD0986 and PD1485 are critical for *Xff* virulence in grapevines. For PD0986, this gene is absent in a *Xf* biocontrol strain EB92-1 that can infect and persist in grapevines but causes only very slight symptoms. When PD0986 is cloned into EB92-1, the transformant induces significantly increased symptoms that are characteristic of PD (Zhang et al., 2015). For PD1485, the knockout mutant was avirulent due to the loss of ability to systemically colonize grapevines (Roper et al., 2007).

Two gene families appeared to have experienced copy number expansion in the *Xt* genome. The first family includes homologs of PD1792 and PD2118, which encode hemagglutinins. These adhesins are antivirulence factors that restrict *in planta* movement by promoting self-aggregation; transposon-insertion mutants of *Xff* PD1792 and PD2118 both exhibit hypervirulence in grapevines (Guilhabert and Kirkpatrick, 2005). Among the representative *Xf* and *Xff* genomes, the median copy numbers of this family are 3 and 8, respectively. In comparison, *Xt* has 12 copies. It remains to be investigated if the copy number variation is linked to protein expression level and virulence. The second family includes a Zot-like toxin (PD0928). Similar to PD0986 (hemagglutinin-like protein), the biocontrol strain EB92-1 lacks the homolog of PD0928 and the transformant that expresses this gene is virulent (Zhang et al., 2015).

## Conclusions

In conclusion, this work reported the complete genome sequence of an important plant-pathogenic bacterium that is endemic to Taiwan. In addition to providing the genomic resource that contributes to the study of this pathogen, this species is the only known sister of *Xf*, which has extensive genetic variations and devastating effects on agriculture worldwide. The availability of this new *Xt* genome sequence provides critical genomic information of a key lineage that may improve the study of *Xylella* evolution and the inference of *Xf* ancestral states. At above-genus level, our genome-scale phylogenetic inference resolved the relationships between *Xylella* and *Xanthomonas*, which are some of the key plant pathogens in the family Xanthomonadaceae.

For gene content analysis, our comparison of the putative virulence genes and pathogenicity factors among representative *Xylella* strains identified the genes that exhibit high levels of conservation or diversity (**Figures 4 and 5**). These genes are promising candidates for future functional studies to investigate the molecular mechanisms of *Xylella* virulence. Previous characterizations of single-gene mutants, particularly those conducted in *Xff*, have provided a strong foundation (Burdman et al., 2011; Rapicavoli et al., 2018). However, it is important to note that the current knowledge of *Xylella* virulence genes is mostly derived from to those strains that are relatively easy to culture and transform, such that only limited diversity has been investigated in molecular genetics studies. Moreover, infection experiments for the investigation of gene functions were limited to a small number of plant species. For further improvements, experimental studies that examine more diverse *Xylella* lineages and plant hosts, as well as the combined effects of multiple virulence genes will be critical.

## Data Availability

The complete genome sequence of *Xylella taiwanensis* PLS229^T^ has been deposited in GenBank/ENA/DDBJ under the accession CP053627. The raw reads have been deposited at the NCBI Sequence Read Archive under the accession numbers SRR11805344 and SRR11805345.

## Funding

The funding was provided by the Ministry of Science and Technology of Taiwan (MOST 106-2923-B-002-005-MY3) to CWT and Academia Sinica to CHK. The funders had no role in study design, data collection and interpretation, or the decision to submit the work for publication.

## Acknowledgments

The Illumina and Oxford Nanopore sequencing library preparation service was provided by the Genomic Technology Core (Institute of Plant and Microbial Biology, Academia Sinica). The Illumina MiSeq sequencing service was provided by the Genomics Core (Institute of Molecular Biology, Academia Sinica).

## Author Contribution

Conceptualization: CWT, CHK

Funding acquisition: CWT, CHK

Investigation: LWW, YCL, CTH

Methodology: CCS, STC, APC, SJC, CHK

Project administration: CWT, CHK

Resources: CCS, CWT, CHK

Supervision: CWT, CHK

Validation: LWW, YCL, CTH

Visualization: LWW, YCL, CTH

Writing – original draft: LWW, CHK

Writing – review & editing: LWW, YCL, CCS, CTH, STC, APC, SJC, CWT, CHK

## Conflict of Interest

The authors declare no conflict of interest.

## Notes

### Competing Interest Statement

The authors have declared no competing interest.

### Summary of Updates

Substantial revision of the "Virulence Genes and Pathogenicity Factors" section in Results and Discussion. Adding new figures to present the results.

